# Transcriptome analysis of mycorrhizal and non-mycorrhizal soybean plantlets upon infection with *Fusarium virguliforme*, one causal agent of sudden death syndrome

**DOI:** 10.1101/388025

**Authors:** N. Marquez, M. L. Giachero, A. Gallou, H. J. Debat, S. Declerck, D. A. Ducasse

## Abstract

Soilborne pathogens represent a threat to agriculture causing important yield losses. The “Sudden Death Syndrome” (SDS), a severe disease in soybean is caused by a complex of *Fusarium* species. This pathosystem has been widely investigated and several strategies were proposed to manage SDS. Although a decrease in symptoms and in the level of root tissue infection particularly by *F. virguliforme* was observed in presence of arbuscular mycorrhizal fungi (AMF), biological control based on AMF has received less attention. Here we report the results, under strict *in vitro* culture experimental conditions, a transcriptional analysis in mycorrhizal versus non-mycorrhizal soybean plantlets upon infection by *F. virguliforme.* An important transcriptional reprogramming was detected following infection by the pathogen. Results revealed 1768 and 967 differentially expressed genes in the AMF-colonized (+AMF+Fv) and non-colonized (−AMF+Fv) plants, respectively. Major transcriptional changes, corresponded to defence response related genes belonging to secondary metabolism, stress and signalling categories. The +AMF+Fv treatment showed the largest number of upregulated genes related to defence, as those encoding for disease resistance proteins, WRKY transcription factors, auxins, receptors kinases, and proteases. Only few genes had primed expression in +AMF+Fv treatment, as those coding for a thaumatin-like protein (TLP) and a pleiotropic drug resistance (PDR) protein. Moreover, +AMF+Fv showed a significant number of downregulated genes related to cell wall modification and peroxidases than – AMF+Fv treatment. This detailed insight increases our knowledge on the transcriptional changes and the potential metabolic pathways involved in the enhanced resistance/tolerance of mycorrhizal plants upon infection with *F. virguliforme.*

## INTRODUCTION

Soybean *(Glycine max* (L.) Merr.) acreage has increased steadily over the past decades, reaching more than 120 million ha in 2017 (FAOSTAT, 2018). It represents nowadays the oilseed crop with the greatest production worldwide. Argentina is the third-largest soybean producer in the world and the second largest in South America after Brazil (FAOSTAT, 2018). Breeding technologies for yield increase, resistance to herbicides or abiotic and biotic stresses, have contributed to expand soybean production (Bilyeu *et al.*, 2016). However, diseases are still a serious threat to this crop. Among the various diseases of soybean (Hartman *et al.*, 2016), sudden death syndrome (SDS) is one of the primary causes of yield losses in South and North American countries (Wrather *et al.*, 2010; Allen, 2017). In North America, SDS is caused by *Fusarium virguliforme*; whereas in South America, it is caused by four *Fusarium* species, *F. virguliforme, F. tucumaniae, F. brasiliense,* and *F. crassistipitatum* (Aoki *et al.*, 2003, 2005, 2012). Symptoms of SDS include root necrosis, tan discoloration of vascular tissues in the lower stem, interveinal leaf chlorosis and necrosis that progresses to premature defoliation (Roy *et al.*, 1997). Several studies have been conducted to evaluate potential control strategies (Wen *et al.*, 2014; S Navi & Rajasab, 2016; Srour *et al.*, 2017; Brzostowski *et al.*, 2018) and it seems that multiple approaches in parallel have to be used for the management of SDS (Hartman *et al.*, 2015). For instance, resistant cultivars in combination with fluopyram seed treatment or in-furrow application have provided some promising results (Kandel *et al.*, 2016).

In the recent years, alternative control measures to pest and diseases have emerged based on the application of nature-based compounds (e.g. plant extracts, microbial metabolites, minerals and ions) (Siah *et al.*, 2018) as well as on the use of bio-control agents (e.g. *Pseudomonas* sp., *Trichoderma* sp. or arbuscular mycorrhizal fungi – AMF), globally termed BCAs (Schouteden *et al.*, 2015; Pandin *et al.*, 2017). This is materialized by a marked increase of biological control products as compared to chemical pesticides, and by an increasing number of products entering the market (Dunham & DunhamTrimmer, 2015).

Arbuscular mycorrhizal fungi are increasingly considered among microorganisms used in biocontrol. These fungi are soil inhabitants forming the most widespread mutualistic symbiosis with plants (Smith & Read, 2008), including many important agricultural crops (Wang & Qiu, 2006). Their effects on plant fitness are largely reported and include a better mineral nutrition and increased ability to overcome biotic and abiotic stresses (Jung *et al.*, 2012; Berruti *et al.*, 2016; Lenoir *et al.*, 2016). For instance, AMF have often been reported to reduce the incidence and/or severity of diseases caused by diverse soilborne fungal pathogens such as *Macrophomina sp., Pythium sp., Fusarium sp., Verticillium sp., Rhizoctonia sp.* (Olawuyi *et al.*, 2014; Srivastava *et al.*, 2014; Eke *et al.*, 2016; Oyewole *et al.*, 2017; Zhang *et al.*, 2018).

The interactions between Soybean and *F. virguliforme* have been widely investigated. In particular molecular approaches (e.g. transcriptional changes in plant roots and pathogen) have received an increasing attention in the recent years (Hartman *et al.*, 2015). For instance, several studies have reported an increased expression of defence related genes in soybean (e.g. phenylpropanoid pathway, signal transduction, transcription factors, and programmed cell death) in response to SDS (Iqbal *et al.*, 2005; Yuan *et al.*, 2008; Radwan *et al.*, 2011). A number of genes encoding enzymes involved in antimicrobial compounds and cell wall degradation have also been identified as putative virulence factors of *F. virguliforme* (Sahu *et al.*, 2017). These studies contributed to expand our understanding of the virulence mechanisms of *F. virguliforme* and the molecular mechanisms deployed by the soybean plant to combat the pathogen. However, to our knowledge, the role of AMF in increasing the resistance/tolerance of soybean to *F. virguliforme* has been little considered to date. Only very recently Giachero *et al.* (2017) proposed the use of *in vitro* cultivation systems for the study of the soybean/AM F/F. *virguliforme* interactions. These systems have been proven to be suitable for studying underground interactions and their associated transcriptional changes (Marquez *et al.*, 2018), since there is no interference with unwanted microbes (Gallou *et al.*, 2010, 2012; Bressano *et al.*, 2010) that could potentially lead to confounding effects (Cameron *et al.*, 2013). Using such system, Giachero *et al.* (2017) observed an attenuation of symptoms in mycorrhizal soybean plants, which was correlated with a reduction in pathogen colonization. This result suggested a mycorrhizal induced protection. However, molecular approaches to the interactions between AMF and *F. virguliforme* on soybean remain fragmentary and argue for further investigation under highly controlled conditions. Therefore, the objective of the present study was to investigate, under *in vitro* culture conditions, transcriptional changes in the roots of mycorrhizal and non-mycorrhizal soybean plantlets upon infection by *F. virguliforme,* increasing our understanding of the potential of these obligate root symbionts as biological control agents against SDS.

## MATERIALS AND METHODS

### Biological Material

Strains of *Rhizophagus irregularis* (Błaszk., Wubet, Renker and Buscot) (Schubler, 2010) as [‘irregulare’] MUCL 41833 and *Fusarium virguliforme* O’Donnell & T. Aoki MUCL 53605 (Aoki *et al.*, 2005) were provided by the Glomeromycota *in vitro* collection and the Mycothèque de l’Université catholique de Louvain (MUCL), respectively. For *R. irregularis,* spores and AMF-colonized root pieces were associated to Ri T-DNA-transformed root organ cultures (ROC) of carrot *(Daucus carota L.* clone DC2) on the modified Strullu-Romand (MSR) medium (Declerck *et al.*, 1998) as detailed in (Cranenbrouck *et al.*, 2005). The cultures were maintained in inverted position in the dark at 27°C for three months. For *F. virguliforme,* a plug of gel containing several macroconidia and mycelium was placed on Potato Dextrose Agar (PDA) and incubated in the dark at 27°C for seven days.

Seeds of *Medicago truncatula* Gaertn. cv. Jemalong A 17 (SARDI, Australia) and soybean *(G. max* (L.) Merr.) cv DON MARIO 4800 were surface-disinfected by immersion in calcium hypochlorite, rinsed three time in deionized sterilized (121 °C for 15 min) water and germinated on Petri plates filled with MSR medium without sucrose and vitamins. The Petri plates were incubated 4 days at 27°C in the dark and then exposed to light (average photosynthetic photon flux (PPF) of 225 μmol m^-2^ s^-1^) for one day.

### Experimental set up

The mycelium donor plant (MDP) *in vitro* culture system developed by Voets *et al.* (2009) for the fast and homogenous AMF colonization of roots of *M. truncatula,* was adapted to soybean in the present study (see for details Giachero *et al.* 2017). Briefly, the cover of a 55 mm diameter Petri plate (named root compartment, RC) was introduced in the base of a 145 mm diameter Petri plate (named hyphal compartment, HC). AMF was associated to roots of *M. truncatula* in the RC and further maintained at 20/18°C (day/night), 70 % relative humidity, with a photoperiod of 16 h d^-1^ and a PPF of 225 μmol m^-2^ s^-1^. A profuse AMF extra-radical mycelium (ERM) network bearing numerous spores extended in the HC of the bi-compartmented Petri plate within a period of 8 weeks. At that time, a 5 days old soybean plantlet was inserted in the HC with the roots in direct contact with the ERM and the shoot extending outside the Petri plate via a hole. In parallel and following the same protocol and timing, soybean plantlets were inserted in culture plates without AMF (i.e. the control treatment).

All the culture plates were incubated for two weeks in a growth chamber under the same conditions as above. At day 7, the RC of the MDP *in vitro* culture systems containing the donor plants were removed from the systems and the empty spaces refilled with 30 ml of fresh MSR medium to allow homogenous regrowth of the ERM. The systems with and without AMF were randomly divided in two groups. Half of the systems were inoculated with *F. virguliforme* using 2 ml suspension of 1.52 × 10^6^ macroconidia ml^−1^ in sterile water. Four treatments were thus considered: AMF-colonized soybean plantlets inoculated (+AMF+Fv) or not (+AMF–Fv) with *F. virguliforme* and non AMF-colonized soybean plantlets inoculated (−AMF+Fv) or not (−AMF–Fv) with *F. virguliforme.* Six biological replicates were set up per treatment. The systems were harvested 72 hours post inoculation (hpi) with *F. virguliforme.* Half of the root material was used to estimate colonization by AMF and/or infection by *F. virguliforme* and the other half was stored at −80°C for RNA extraction.

### Assessment of AMF root colonization and root infection by *F. virguliforme*

Root colonization by AMF and infection by *F. virguliforme* were assessed at harvest (i.e., 72 hpi with the pathogen). Fresh roots were cleared in 10% KOH at room temperature for 3 h, rinsed with distilled water, bleached and acidified with HCl and stained with Trypan blue at room temperature for 15 min (Phillips & Hayman, 1970).

For AMF, root fragments (1 cm long) were mounted on microscope slides, and 200 intersections were observed under a dissecting microscope (Olympus BH2, Olympus Optical, GmbH, Germany) at 10-40 x magnifications. Total root colonization (% RC), abundance of arbuscules (% A) and intraradical spores/vesicles (% V) were determined according to McGonigle *et al.* (1990). Infection by *F. virguliforme* was estimated following the methodology described by McGonigle *et al.* (1990) slightly modified as decribed by Giachero *et al.* (2017). For all the replicates, between 2 and 4 slides, each covered with 10 root fragments (10 mm long), were evaluated. Intersections were counted under the microscope at 10-40 x magnifications as ‘absence’ or ‘presence’ of hyphae, arbuscules or vesicles/spores of the AMF and presence of hyphae of the pathogen. AMF and *F. virguliforme* mycelia were clearly distinguished by their morphology and growing pattern in the +AMF+Fv treatment. The percentage of infected roots was estimated as the ratio between infected root pieces and total number of root pieces examined. Moreover, symptoms caused by *F. virguliforme* on the entire root system was qualitatively evaluated, under binocular microscope (Olympus SZ61) at 5 x magnification.

### RNA extraction and cDNA

Total RNA was extracted with the RNeasy Plant Mini kit (Qiagen, Valencia, USA) and treated afterward with the TURBO DNA-free^TM^ kit (Ambion, Austin, USA) according to the manufacturer’s instructions. Concentration and purity of total RNA were determined in a NanoDrop^®^-ND 1000 UV-vis Spectrophotometer (NanoDrop Technologies, Wilmington, USA), and the total RNA quality was tested using the Agilent 2100 BioAnalyzer as recommended by the manufacturer’s protocol (RNA 6000 Nano Assay Protocol, Agilent Technologies, Santa Clara, USA). For each treatment, total RNA of two replicates was randomly selected and pooled (i.e., biological replicate). Thus, three independent biological replicates were used per treatment for the microarray experiment.

### Microarray experiment

A two-colour microarray-based gene expression analysis protocol was used. cDNA labelling, microarray hybridization and pre-processing of microarray data was done as described in Márquez *et al.* (2018).

The oligonucleotide arrays (Agilent-016772 G. max Oligo Microarray Agilent 4×44K) were hybridized, stained, washed and scanned at the Platform of Applied Molecular Technologies (https://uclouvain.be/fr/instituts-recherche/irec/ctma) of the Université catholique de Louvain (UCL, Belgium). All treatments replicates (including −AMF–Fv) were hybridized with a pool of the three biological replicates of −AMF–Fv treatment (competitive hybridization).

Microarray images were imported in the Agilent Feature Extraction (FE) software (version 9.1.3.1) and aligned with the appropriate array grid template file (Agilent Technologies, Santa Clara, USA).

Finally, for genes expression, selected comparison among treatments was performed by appropriate contrasts (+AMF–Fv vs −AMF–Fv; – AMF+Fv vs – AMF–Fv; +AMF+Fv vs −AMF–Fv; +AMF+Fv vs −AMF+Fv). Genes were considered as differentially regulated if p value was below or equal to 0.005 according to Di Rienzo, Guzman, and Casanoves (DGC) test (Di Rienzo *et al.*, 2002) and a three-fold cut-off.

Microarray data have been deposited in Gene Expression Omnibus (GEO-NCBI) and are accessible through GEO Series accession number GSE108964.

### Gene Ontology (GO) and Functional Annotation

For Gene Ontology (GO) analysis “Mercator Automated Sequence Annotation Pipeline” and “MapMan” were used to generate functional assignments for each input gene and data visualization/interpretation of soybean gene expression involved in biotic stress.

Functional annotation of differentially expressed genes was generated with the KAAS tool (KEGG Automatic Annotation Server) as implemented in https://www.genome.jp/kegg/kaas/resulting in KO (KEGG Orthology) assignments and automatically generated KEGG pathways (Moriya et al. 2007). The obtained KO terms were traced to the corresponding soybean ESTs and integrated with their corresponding microarray FC values.

### Statistical analysis

Percentages of root colonization by AMF or root infection by *F. virguliforme* were analysed using one-way analysis of variance (ANOVA). Data were subjected to the LSD Fisher’s honest significant difference (HSD) test in order to identify the significant differences (p ≤ 0.05) between treatments.

Gene expression data (i.e. relative expression ratio) was analysed by two-way ANOVA. DGC test model was conducted to identify significant differences (p ≤ 0.05) between the treatments (Di Rienzo *et al.*, 2002). One-way and two-way ANOVA were performed using the software InfoStat (Di Rienzo *et al.*, 2013).

## RESULTS

### Root colonization by *R. intraradices*

At the end of the experiment, plants were harvested and root AMF colonization evaluated. The %RC was 36.2 ± 12.8 and 42.9 ± 14.0 in the +AMF–Fv and +AMF+Fv treatments, respectively. The %A was 18.5 ± 10.5 and 16.2 ± 4.4 in the +AMF–Fv and +AMF+Fv treatments, respectively. The %V was 2.2 ± 2.3 and 1.5 ± 2.4 in the +AMF–Fv and +AMF+Fv treatments, respectively. None of these parameters significantly differed between the two treatments according to LSD Fisher test (p ≤ 0.05). No AMF root colonization was noticed in the −AMF+Fv and −AMF–Fv treatments.

### Root infection by *F. virguliforme*

A fast and dense development of hyphae was clearly visible on the surface of the MSR medium 72 hpi with *F. virguliforme.* Root penetration and subsequent tissues invasion was confirmed by microscopic observation. Root infection by *F. virguliforme* was significantly higher according to LSD Fisher test (p ≤ 0.05) in the −AMF+Fv treatment (i.e. 44.0% ± 2.9) as compared to the +AMF+Fv treatment (i.e. 35.7% ± 1.7). Moreover, 72 hpi with *F. virguliforme,* necrotic areas were observed on the surface of the roots of non-mycorrhizal and mycorrhizal soybean plantlets, while no symptoms were detected in roots of +AMF–Fv and −AMF–Fv treatment.

### Differential expression of genes

Microarray experiments were performed to profile soybean gene expression during AMF colonization and *F. virguliforme* infection. We used the term ‘gene’ for a probe set representing a given transcript.

The total number of differentially expressed genes in the three treatments was shown in a Venn diagram (Figure 1). Soybean roots revealed 1066 and 702 genes up and down-regulated, respectively in the +AMF+Fv treatment, 673 and 294 up and down-regulated genes, respectively in the −AMF+Fv treatment and 152 and 23 up and down-regulated genes, respectively in the +AMF–Fv treatment. As shown in Figure 1, 45 genes were co-regulated under the three treatments. Following *F. virguliforme* inoculation, 797 genes were co-regulated in the −AMF+Fv and +AMF+Fv treatments. Furthermore, following mycorrhizal colonization, 124 genes were co-regulated between +AMF–Fv and +AMF+Fv treatments. Finally, 892 differentially expressed genes were unique to the +AMF+Fv treatment.

**Figure 1:**
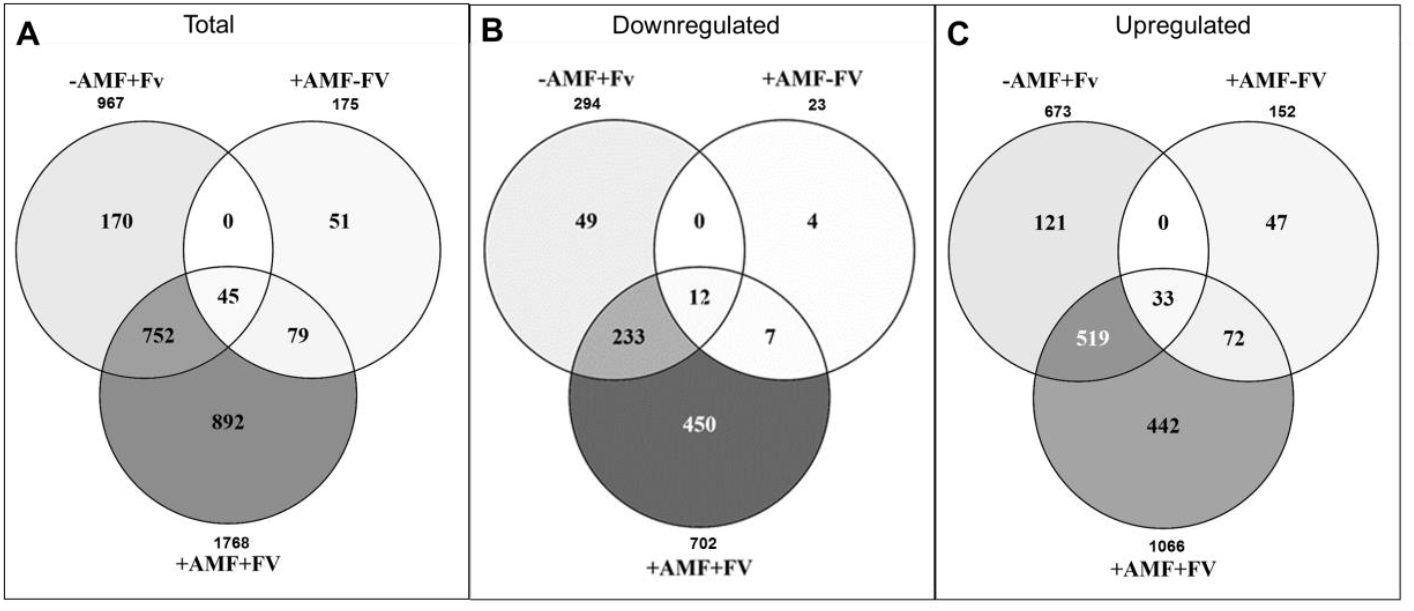
Venn diagram showing the relationships between the total (A), downregulated (B) and upregulated (C) differentially expressed genes in roots of pre-mycorrhizal plantlets infected (+AMF+Fv) or not (+AMF–Fv) with *Fusarium virguliforme* and in roots of non-mycorrhizal plantlets infected with *F. virguliforme* (−AMF+Fv). Genes were considered as differentially regulated if p ≤0.005 (DGC comparative method) and the values of fold change compared with the −AMF–Fv treatment was ≥3.0 or ≤3.

### Functional assessment of differentially expressed genes

MapMan software was used for data visualization and interpretation of soybean genes expression. Differentially expressed genes were assigned into the different metabolic pathways (Figure 2). Results revealed that 50% of the differentially regulated genes in each treatment could not be assigned to a specific function or are miscellaneous (Figure 2A). Among the other 50%, major transcriptional changes were observed in functions related to RNA, signalling, transport, proteins, cell wall, stress, secondary metabolism, and hormone metabolism.

**Figure 2:**
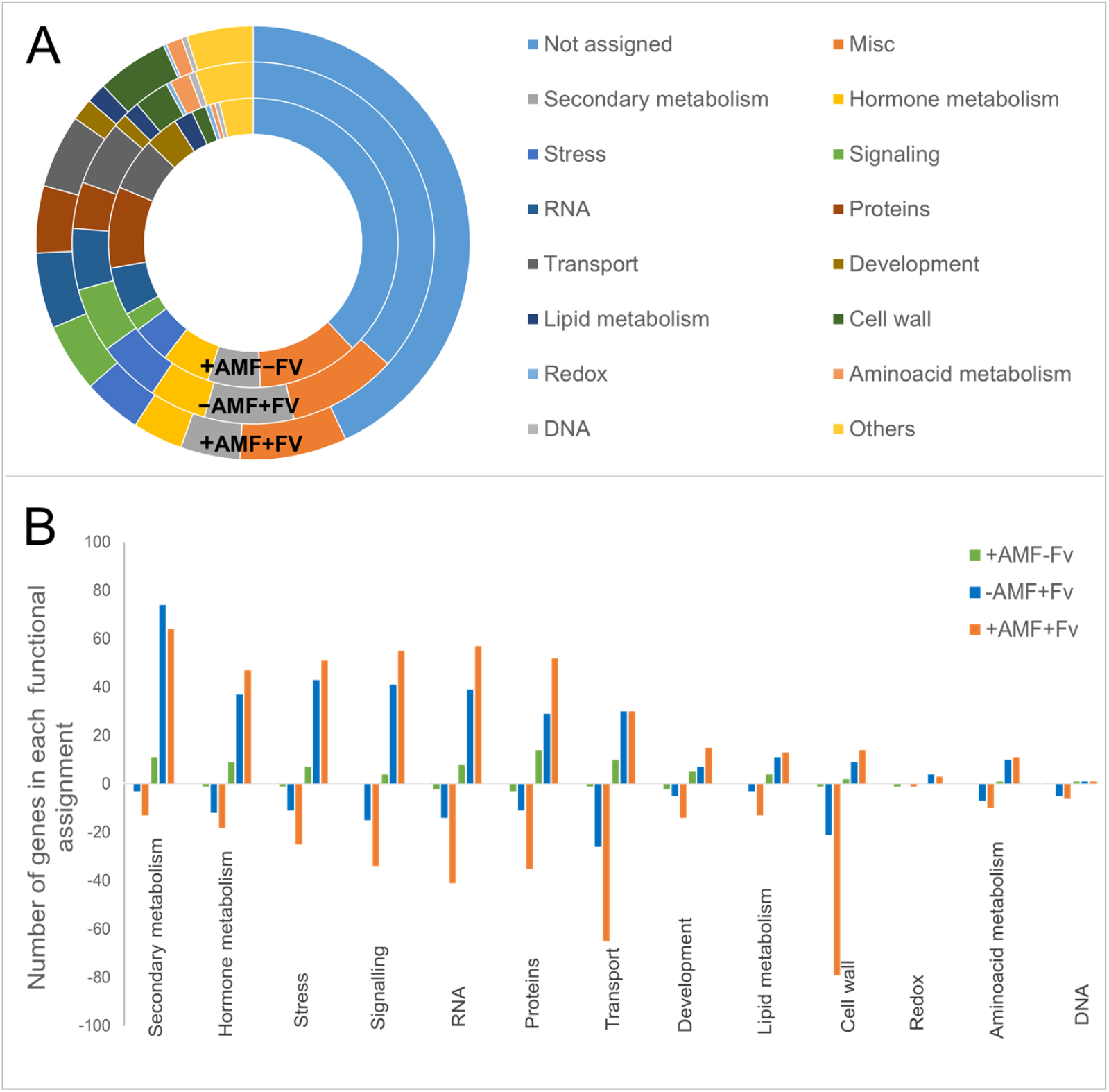
Functional assignment determined by the MapMan software of genes differentially regulated (A), upregulates and downregulated (B) in roots of pre-mycorrhizal plantlets infected (+AMF+Fv) or not (+AMF–Fv) with *Fusarium virguliforme* and in roots of non-mycorrhizal plantlets infected with *F. virguliforme* (−AMF+Fv). Genes were considered as differentially regulated if p ≤0.005 (DGC comparative method) and the values of fold change compared with the −AMF–Fv treatment was ≥3.0 or ≤3.

After soybean inoculation with *F. virguliforme* an important transcriptional reprogramming was detected (Figure 2B). Results showed that the major changes corresponded to the defence response related genes belonging to secondary metabolism, stress and signalling categories. However, roots of pre-mycorrhizal plantlets (+AMF+Fv) showed larger number of differentially expressed genes in most of the categories as compared to non-mycorrhizal plantlets (−AMF+Fv).

Moreover, a considerable number of transport, cell wall and signalling related genes were down-regulated in presence of the pathogen. Noteworthy, the number of downregulated genes were remarkably greater in the +AMF+Fv treatment.

### Transcriptional changes of biotic stress related genes

With the purpose to understand the plant defence response, the transcriptional changes were analysed with special focus on biotic stress-related genes in the +AMF+Fv and −AMF+Fv treatments. MapMan biotic stress graph (Figure 3) revealed the up-regulation of numerous genes. Most of them were related to signalling, such as receptor kinases (DUF26, leucine rich repeat, MAPK, wall associated kinase and S-locus glycoprotein like) and secondary metabolism related genes associated with simple phenols, flavonoids, and lignin biosynthesis. Genes encoding a ferulate 5-hydroxylase (F5H), Cytochrome P450 71D8 (CCoAOMT), and glycosyltransferase were notably upregulated. Moreover, an important number of hormone signalling related genes (auxins, ABA, ethylene, jasmonic acid, and salicylic acid), pathogenesis-related proteins (PR1 and PR5), dirigent-like proteins, glutathione S transferases, and transcription factors (WRKY, MYB, and ERF) were over-expressed. The 948 co-regulated genes between +AMF+Fv and −AMF+Fv treatments did not show significant differences in their fold change. Only a few genes had primed expression in +AMF+Fv treatment, as those coding for a benzyl alcohol O-benzoyltransferase-like, an auxin-induced protein, a RNA-binding protein ARP1, pleiotropic drug resistance (PDR) protein or a protein NRT1 (Table 1). Nevertheless, the +AMF+Fv treatment showed major number of upregulated genes related to defence such as those encoding for disease resistance proteins, WRKY transcription factors, auxins, receptors kinases, and proteases (Supplementary Table 1).

**Figure 3:**
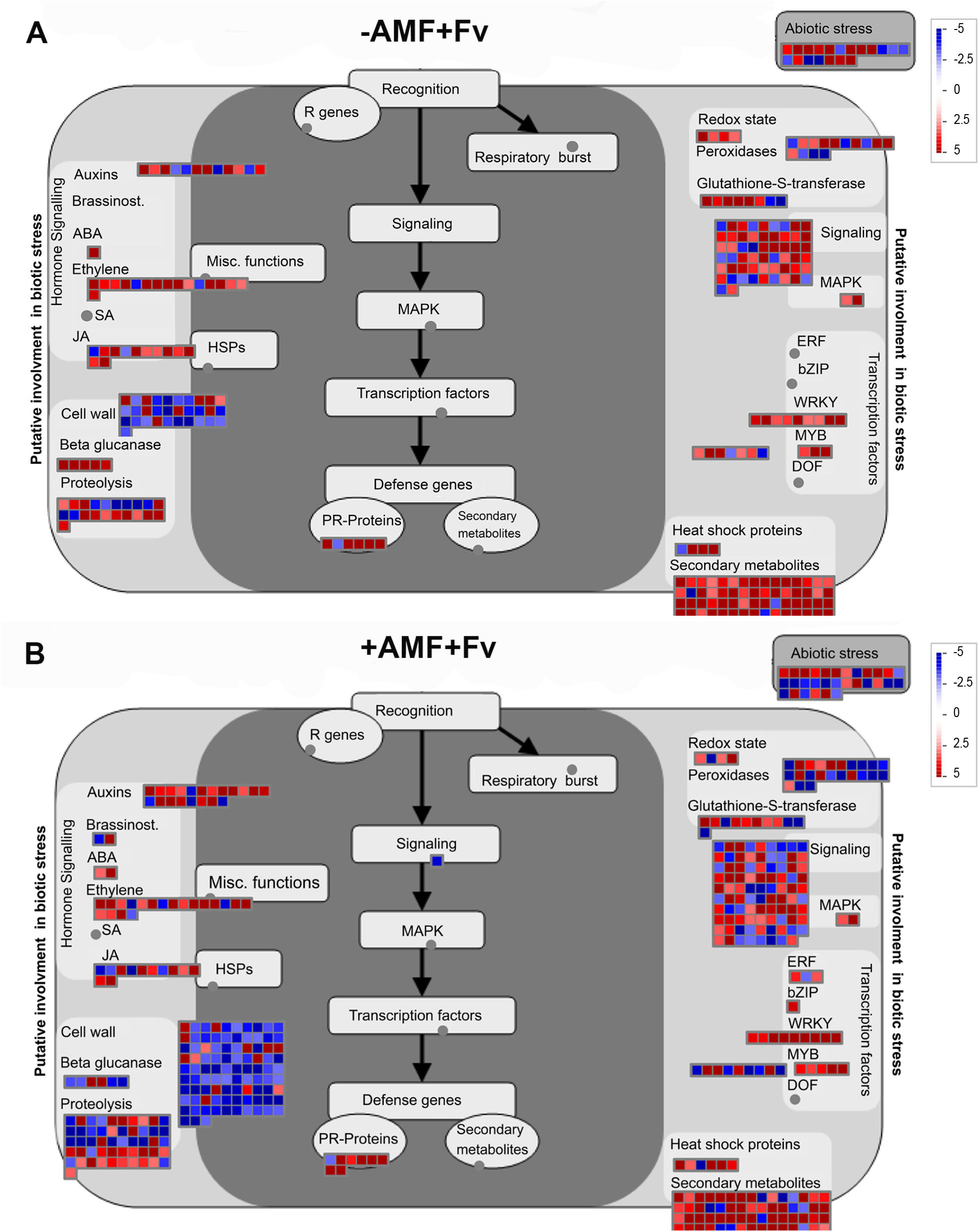
Transcriptional changes in soybean roots after AMF pre-colonization and *Fusarium virguliforme* infection. (A) MapMan Biotic Stress graph generated using the 1740 distinctively regulated genes identified in roots 72 hours post inoculation (hpi) with *F.virguliforme* versus the non-mycorrhizal soybean plantlets not infected with *F*. *virguliforme* (B) MapMan Biotic Stress graph generated using the 1398 genes distinctively regulated identified in roots of AMF colonized plantlets 72 hpi with *F.virguliforme* versus the non-mycorrhizal soybean plantlets not infected with *F.virguliforme.* The fold change is displayed as illustrated in the colour bar of the panel (blue is downregulated and red is upregulated).

**Table 1:** List of genes with primed expression in mycorrhizal soybean roots infected (+AMF+Fv) or not (+AMF–Fv) with *Fusarium virguliforme* and non-mycorrhizal soybean roots infected with *F. virguliforme* (−AMF+Fv). Genes were considered as differentially regulated if p ≤0.005 and the values of fold change compared with the −AMF–Fv treatment was ≥3.0 or ≤3. Data is organized by GeneBank accession (ID).

| ID | Description | -AMF+Fv |  | +AMF+Fv |  | +AMF-Fv |  |
| --- | --- | --- | --- | --- | --- | --- | --- |
|  |  | FC | p-value | FC | p-value | FC | p-value |
| AI443411.1 | Glycine max DNA-damage-repair/toleration protein DRT100 | 2,08 | 0,06 | 6,72 | 0,00 | 1,63 | 0,15 |
| AW185044.1 | Glycine max glycolipid transfer protein 3-like | 3,15 | 0,01 | 14,73 | 0,00 | 4,73 | 0,00 |
| AW278861.1 | Glycine max cytochrome P450 71D9-like | 2,94 | 0,07 | 10,46 | 0,00 | -1,08 | 0,88 |
| AW348060.1 | Glycine max auxin-responsive protein IAA14-like | 1,03 | 0,89 | 3,49 | 0,00 | 1,09 | 0,69 |
| AW348589.1 | Glycine max protein NRT1/ PTR FAMILY 3.1 | 7,52 | 0,00 | 29,75 | 0,00 | 2,43 | 0,05 |
| AW349280.1 | Glycine max xyloglucan endotransglucosylase/hydrolase protein 31-like | 6,23 | 0,01 | 19,62 | 0,00 | 1,52 | 0,42 |
| BE210309.1 | Glycine max arabinogalactan peptide 14 | 1,26 | 0,52 | 6,24 | 0,00 | -1,01 | 0,99 |
| BE346184.1 | Glycine max polygalacturonase At1g48100-like | 1,16 | 0,63 | 3,59 | 0,00 | 1,16 | 0,60 |
| BE555205.1 | Glycine max vegetative cell wall protein gp1-like | 1,07 | 0,69 | 4,03 | 0,00 | 1,29 | 0,14 |
| BE821916.1 | Glycine max transcription factor MYB86 | 1,64 | 0,24 | 6,60 | 0,00 | 1,70 | 0,17 |
| BE822069.1 | Unknown | 3,16 | 0,08 | 13,78 | 0,00 | 2,63 | 0,11 |
| BE822496.1 | Glycine max transcription factor bHLH25-like | 1,21 | 0,39 | 4,60 | 0,00 | 1,31 | 0,21 |
| BU546292.1 | Glycine max pleiotropic drug resistance protein 3 | 1,38 | 0,37 | 7,04 | 0,00 | 2,32 | 0,02 |
| BU551023.1 | Glycine max membrane protein of ER body 2-like | 1,10 | 0,81 | 5,98 | 0,00 | 1,55 | 0,25 |
| BW651899.1 | Glycine max auxin-induced protein | 8,50 | 0,01 | 31,68 | 0,00 | 3,34 | 0,06 |
| BW654381.1 | Glycine max protein GRIM REAPER-like | 1,54 | 0,31 | 5,01 | 0,00 | 1,39 | 0,39 |
| BW654899.1 | Glycine max thiamine thiazole synthase 2, chloroplastic-like | 3,60 | 0,08 | 14,37 | 0,00 | 2,31 | 0,19 |
| BW661061.1 | Glycine max nicotianamine synthase-like | 1,81 | 0,27 | 7,83 | 0,00 | 1,43 | 0,46 |
| BW668263.1 | Unknown | 3,97 | 0,01 | 13,93 | 0,00 | 1,68 | 0,22 |
| BW670549.1 | Glycine max cultivar BX10 extensin-like protein mRNA, complete cds | -1,06 | 0,90 | 10,87 | 0,00 | 1,77 | 0,19 |
| CA853372.1 | Glycine max peroxidase 3-like | 1,03 | 0,92 | 3,69 | 0,00 | 1,08 | 0,80 |
| CF921629.1 | Glycine max germin-like protein 9 mRNA, complete cds | 1,51 | 0,47 | 10,07 | 0,00 | 1,22 | 0,70 |
| CO983452.1 | Glycine max polygalacturonase At1g48100 | 2,23 | 0,16 | 10,40 | 0,00 | 1,67 | 0,30 |
| CX547879.1 | Glycine max probable xyloglucan endotransglucosylase/hydrolase protein 33 | 1,68 | 0,23 | 6,09 | 0,00 | 1,21 | 0,62 |
| CX547880.1 | Glycine max thaumatin-like protein 1b | 1,53 | 0,29 | 5,44 | 0,00 | 1,73 | 0,14 |
| CX703081.1 | Glycine max codeine O-demethylase-like | 2,83 | 0,00 | 8,61 | 0,00 | 2,22 | 0,01 |
| CX703135.1 | Glycine max organic cation/carnitine transporter 3-like | 2,54 | 0,02 | 10,53 | 0,00 | 1,46 | 0,24 |
| CX704036.1 | Glycine max benzyl alcohol O-benzoyltransferase-like | 13,34 | 0,01 | 60,20 | 0,00 | 4,90 | 0,04 |
| CX706218.1 | Glycine max probable RNA-binding protein ARP1 | 6,40 | 0,00 | 28,22 | 0,00 | 3,77 | 0,01 |

Regarding downregulated genes upon *F. virguliforme* infection, biotic stress graph revealed a strong down-regulation of cell wall associated genes encoding xyloglucan endotransglycosylase, invertases, pectin lyase-like protein, and other esterases. Mycorrhizal plants (+AMF+Fv) showed major number of downregulated genes, encoding cell wall modification related proteins and peroxidases, than −AMF+Fv treatment.

Finally, a set of genes related to biotic stress was observed to be expressed exclusively in mycorrhizal plants. Those co-regulated genes between +AMF+Fv and +AMF–Fv treatments were related to gibberellin (GA) signalling, pathogen related proteins (PRP) (PR1 and PR5), serine proteases and lectin precursors (Soybean agglutinins-SBA) (Supplementary Table 1).

### KEGG pathway enrichment analysis

The sequences reported for the genes with differential expression were mapped in the database resource Kyoto Encyclopedia Genes and Genomes (KEGG) for a better understanding of the defence response during the triple interaction soybean/AMF/pathogen. Phenylpropanoid biosynthesis was the metabolic pathway where the highest number of differentially expressed sequences was found (Figure 4A and 4B). This pathway showed significant changes of genes encoding phenylalanine ammonia-lyase, 4-coumarate-CoA ligase, shikimate O-hydroxycinnamoyltransferase, caffeoyl-CoA O-methyltransferase, cinnamyl-alcohol dehydrogenase, beta-glucosidase, peroxidase and coniferyl-alcohol glucosyltransferase upon *F. virguliforme* infection (Figure 4B), while AMF colonization (+AMF–Fv) did not caused major changes. Moreover, +AMF+Fv and−AMF+Fv treatments showed 27 (19 dowregulated and 8 upregulated) and 18 (9 dowregulated and 9 upregulated) differentially expressed genes respectively, encoding a peroxidase enzyme [EC: 1.11.1.7]. Remarkably, pre-mycorrhizal soybean plants led to a robust downregulation of peroxidase genes in presence of the pathogen (Figure 5).

**Figure 4:**
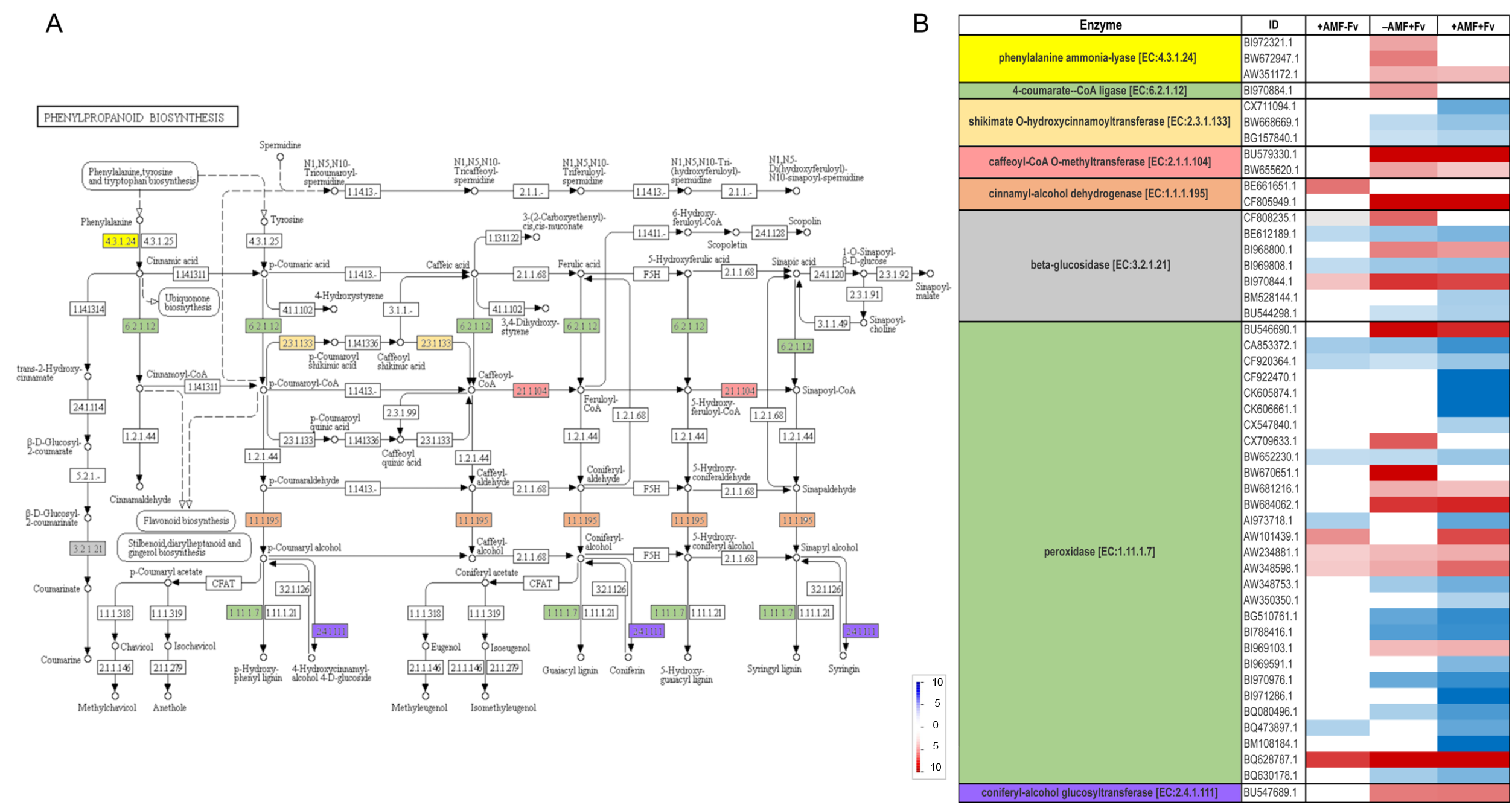
Image obtained through KEGG Pathway Maps showing the phenylpropanoid biosynthesis scheme (A). Differentially expressed genes encoding enzymes from the phenylpropanoids pathway are shown with different colors and codes. Image credits: Kanehisa Laboratories, Japan. (B) Heat maps indicates the fold change (FC) of phenylpropanoid biosynthesis related genes in mycorrhizal soybean roots infected (+AMF+Fv) or not (+AMF–Fv) with *Fusarium virguliforme* and non-mycorrhizal soybean roots infected with *F. virguliforme* (−AMF+Fv) after AMF pre-colonization and *Fusarium virguliforme* infection. Genes were considered as differentially regulated if p 0.005 (DGC comparative method) and the values of fold change compared with the −AMF-Fv treatment was ≥3.0 or ≤3. The fold change is displayed as illustrated in the colour bar of the panel (blue is downregulated and red is upregulated).

**Figure 5:**
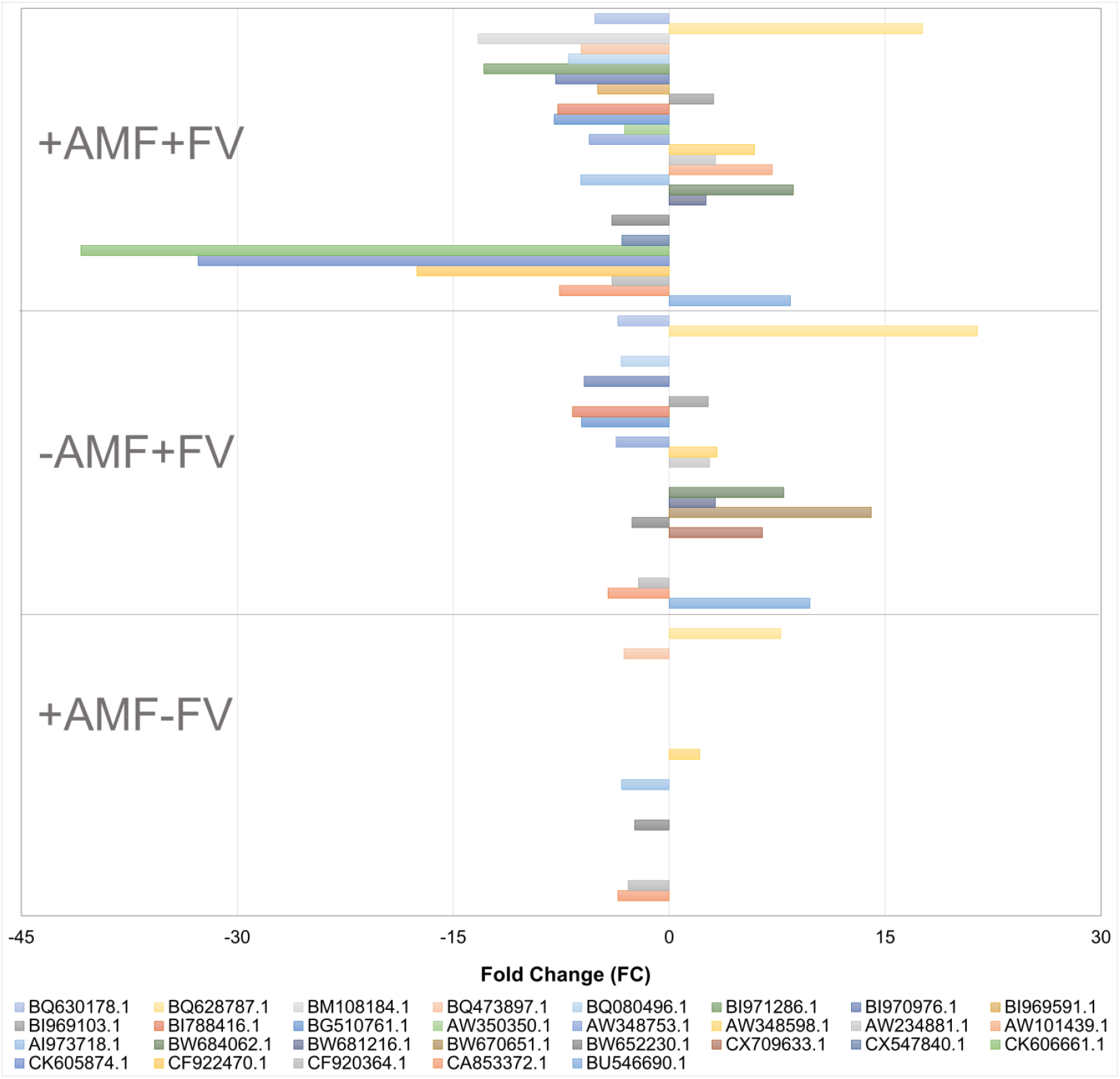
Fold change (FC) of peroxidase related genes in mycorrhizal soybean roots infected (+AMF+Fv) or not (+AMF–Fv) with *Fusarium virguliforme* and non-mycorrhizal soybean roots infected with *F. virguliforme* (−AMF+Fv) after AMF pre-colonization and *Fusarium virguliforme* infection. Genes were considered as differentially regulated if p ≤0.005 (DGC comparative method) and the values of fold change compared with the −AMF–Fv treatment was ≥3.0 or ≤3.

## DISCUSSION

The use of AMF for helping plants fend off pests and diseases is considered a promising alternative or complementary approach to the application of chemical pesticides (Al-Askar & Rashad, 2010; Duhamel & Vandenkoornhuyse, 2013; Nadeem *et al.*, 2014; Dar & Reshi, 2017). Bioprotection with AMF has been amply described in different plant systems. Various mechanisms (e.g. improved plant nutrition, competition) have been proposed (Azcón-Aguilar & Barea, 1997; Schouteden *et al.*, 2015) with an increasing attention in the last decennia to a modulation of plant defence responses upon establishment of the AMF, resulting in a primed state of the plant that lead to a more efficient activation of defence mechanisms following attack by enemies (Pozo *et al.*, 2009; Jung *et al.*, 2012; Cameron *et al.*, 2013). However, only a limited proportion of studies has been conducted under strict *in vitro* culture conditions (Cameron *et al.*, 2013), thus in the absence of any potential confounding effects caused by unwanted microbial contaminants or environmental factors that may impact e.g. plant defence responses. For instance Giachero *et al.* (2017) using an *in vitro* cultivation system, demonstrated a decrease in symptoms and in the level of root tissues infection in AMF-colonized soybean plants attacked by *F. virguliforme.* Using the same system, Marquez *et al.* (2018), investigated transcriptional changes in mycorrhizal and non-mycorrhizal soybean plants upon infection with the fungal pathogen *Macrophomina phaseolina* revealing the complex underlying mechanisms involved in AMF-mediated biocontrol. Transcriptional changes during interaction between soybean plants and *F. virguliforme* have already been analysed and contributed to increasing our understanding of the virulence mechanisms of *F. virguliforme* to produce SDS (Chang *et al.*, 2016; Sahu *et al.*, 2017) and the molecular response of the plant to combat the pathogen (Radwan *et al.*, 2011; Ngaki *et al.*, 2016). Likewise, several studies have focused on the transcriptome profiles of AMF-colonized plants with the aim of analysing wide-scale gene reprogramming during the establishment and development of arbuscular mycorrhizal symbioses (Liu *et al.*, 2007; Gallou *et al.*, 2012) or during interaction of AMF-colonized plants with below-(e.g. Marquez *et al.*, 2018) and above-ground (Gallou *et al.*, 2011) pathogens. Accordingly, the aim of this work was to analyse the soybean trancriptome of AMF-colonized and non-colonized plants in the early stages of attack by *F. virguliforme.*

Earlier results identified significant transcriptional changes in soybean plants in response to *F. virguliforme.* Most of the differentially expressed genes were upregulated and those with assigned function encoded signalling, defence, and secondary metabolism related proteins (Radwan *et al.*, 2011; Ngaki *et al.*, 2016). However, further studies are still required to reveal the role of a large number of differentially expressed genes encoding proteins of no-assigned or unknown functions that are not considered and could be involved in plant immunity. In the present study, an increased transcription of genes encoding PRP, serine proteases, receptor kinases, and phenylpropanoid pathway derivatives were observed in the AMF-colonized soybean roots. Moreover, significant transcriptional changes were noticed in mycorrhizal and non-mycorrhizal soybean plants upon infection by *F. virguliforme.* Microarray analysis 72 hpi with *F. virguliforme* revealed that many changes in gene expression were common between mycorrhizal and non-mycorrhizal colonized roots, while others were unique in the AMF-colonized plants infected by the pathogen. Most of the 797 co-regulated genes were related to signalling, secondary metabolism, hormone signalling, pathogenesis-related proteins, and transcription factors. Noticeably, soybean plants were going through a defence response after *F. virguliforme* infection, via increased expression of defence related genes. However, AMF-colonized plants showed higher number of defence expressed genes as compared to non AMF-colonized plants. Part of these induced genes had an enhanced expression in the mycorrhizal plants infected by the pathogen; a phenomenon termed priming (Conrath *et al.*, 2002, 2015; Conrath, 2006); Priming of defence response was proposed to mediate the induced resistance, resulting in the pre-conditioning of the tissues for efficient activation of plant defences upon a challenger attack (Pozo & Azcón-Aguilar, 2007; Pieterse *et al.*, 2014).

Thaumatin-like protein (TLPs) gene was primed in mycorrhizal soybean plants after *F. virguliforme* infection. Thaumatin-like proteins are a group of PRP (PR-5) that are induced in plants in response to infection by pathogens, elicitors and/or stress factors (Liu *et al.*, 2010; Rather *et al.*, 2015). Most of PR-5 genes are believed to have an antifungal activity, because of their enzymatic activities able of degrading fungi cell walls (Vigers *et al.*, 1992; Ho *et al.*, 2007). It has been reported that PR-5 over-expression enhances tolerance in several host-pathogens interactions (Acharya & Pal, 2012; Singh *et al.*, 2013). Nevertheless, Szwacka et al. (2002) did not found a relationship between thaumatin-protein accumulation in transgenic plants and the increased tolerance phenotype. The present study also showed an enhanced expression of a pleiotropic drug resistance (PDR) protein in the mycorrhizal plants infected by the pathogen. The PDR transporters are expressed in various plant biological processes, including response against biotic and abiotic stresses and the transport of a diverse array of molecules across membranes (Crouzet *et al.*, 2013; Nuruzzaman *et al.*, 2014; Migocka *et al.*, 2017; Xu *et al.*, 2018). PDR-like genes have been reported to confer resistance against *Alternaria alternata* in tobacco plants (Xu *et al.*, 2018). Hence, it would be interesting to elucidate the function of this PDR transporters gene family in the soybean-F. *virguliforme* interaction and their potential role in plant defence.

Additionally, the transcriptional reprogramming during AMF establishment (Garcia-Garrido *et al.*, 2002; Liu *et al.*, 2007; López-Ráez *et al.*, 2010) could help the plant to recognize and resist more efficiently a pathogen attack. Lectin receptor kinases (LecRKs), which in this study were induced exclusively in mycorrhizal plants, are supposed to play important roles in sensing alterations at the plant cell wall and, subsequently, to mediate response reactions (Frenzel *et al.*, 2006; Bouwmeester *et al.*, 2011, 2014; Nivedita *et al.*, 2017; Wang & Bouwmeester, 2017). Moreover, receptor kinases are proposed to positively regulate plant immunity at the transcriptional level and to mediate priming regulation (Singh *et al.*, 2012; Woo *et al.*, 2016). Thus, the enhanced resistance/tolerance of mycorrhizal soybean plants to *F. virguliforme* could not only be a consequence of the defence priming.

Phenylpropanoid biosynthesis was the metabolic pathway showing the highest number of differentially expressed genes in the mycorrhizal and non-mycorrhizal soybean plants upon *F. virguliforme* infection. This pathway has a wide variety of intermediates both as structural and signalling molecules (Cheynier *et al.*, 2013). Several genes encoding eight different enzymes in this pathway were affected upon pathogen infection. Particularly, in the mycorrhizal soybean plants a strong downregulation of peroxidase genes was noticed in presence of the pathogen. The diverse peroxidase activities facilitate antagonistic reactions in plants, such as generation/scavenging of ROS and loosening/stiffening of the cell wall (Shigeto & Tsutsumi, 2016). Each peroxidase function is likely to depend on the spatiotemporal regulation of each peroxidase gene expression, and the substrate selectivity of peroxidase. In this context, and being aware of the large number of peroxidase isoforms in a single plant species, it is difficult to determine their role in the different biological processes and especially in plant defence response. However, the downregulation of peroxidase expression noticed in the mycorrhizal plants in presence of the pathogen suggest that it may be part of a signalling mechanism for an induced response, possibly linked with the attenuation of SDS symptoms in the AMF-colonized plants.

The results here reported demonstrate the complexity of the net that dynamically articulates the interactions between plant and two microbes, one beneficial and the other pathogenic. Although the oversimplification of the experimental design is intended for clarifying the results, some of the results will require further experiments to deeply explain them. Nevertheless, the detailed insight provided by this study contributes to increase our understanding of the complex interaction between mycorrhizal soybean and *F.virguliforme* and of the important role of AMF in the defence response regulation against this soilborne pathogen. AMF, as other plant associated beneficial microbes, is called to play a key role as an ecological engineer to solve environmental stress problems (Kumar & Verma, 2018).

## ACKNOWLEDGEMENTS

The authors would like to thank ANPCyT and CONICET doctoral fellowship (N. M.).

---

**Supplementary Table 1:** List of differentially expressed genes in mycorrhizal soybean roots infected (+AMF+Fv) or not (+AMF–Fv) with *Fusarium virguliforme* and non-mycorrhizal soybean roots infected with *F. virguliforme* (−AMF+Fv). Genes were considered as differentially regulated if p ≤0.005 and the values of fold change compared with the −AMF–Fv treatment was ≥3.0 or ≤3.Data is organized by GeneBank accession (ID).

## REFERENCES

Acharya K, Pal AK, 2012. Overexpression of Camellia sinensis Thaumatin-Like Protein, CsTLP in Potato Confers Enhanced Resistance to Macrophomina phaseolina and Phytophthora infestans Infection.

Al-Askar a. a. A, Rashad YMM, 2010. Arbuscular mycorrhizal fungi: A biocontrol agent against common bean Fusarium root rot disease. Plant Pathology Journal 9, 31–38.

Allen T and others, 2017. Soybean Yield Loss Estimates Due to Diseases in the United States and Ontario, Canada, from 2010 to 2014. Plant Health Progress.

Aoki T, O’Donnell K, Homma Y, Lattanzi AR, 2003. Sudden-death syndrome of soybean is caused by two morphologically and phylogenetically distinct species within the Fusarium solani species complex F. virguliforme in North America and F. tucumaniae in South America. Mycologia 95, 660–684.

Aoki T, O’Donnell K, Scandiani MM, 2005. Sudden death syndrome of soybean in South America is caused by four species of Fusarium: Fusarium brasiliense sp. nov., F. cuneirostrum sp. nov., F. tucumaniae, and F. virguliforme. Mycoscience 46, 162–183.

Aoki T, Scandiani MM, O’Donnell K, 2012. Phenotypic, molecular phylogenetic, and pathogenetic characterization of Fusarium crassistipitatum sp. nov., a novel soybean sudden death syndrome pathogen from Argentina and Brazil. Mycoscience 53, 167–186.

Azcón-Aguilar C, Barea JM, 1997. Arbuscular mycorrhizas and biological control of soil-borne plant pathogens--an overview of the mechanisms involved. Mycorrhiza 6, 457–464.

Berruti A, Lumini E, Balestrini R, Bianciotto V, 2016. Arbuscular mycorrhizal fungi as natural biofertilizers: Let’s benefit from past successes. Frontiers in Microbiology 6, 1–13.

Bilyeu K, Ratnaparkhe MB, Kole C, 2016. Genetics, Genomics, and Breeding of Soybean. CRC Press.

Bouwmeester K, Han M, Blanco-Portales R et al., 2014. The Arabidopsis lectin receptor kinase LecRK-I.9 enhances resistance to Phytophthora infestans in Solanaceous plants. Plant Biotechnology Journal 12, 10–16.

Bouwmeester K, de Sain M, Weide R et al., 2011. The lectin receptor kinase LecRK-I.9 is a novel Phytophthora resistance component and a potential host target for a RXLR effector. PLoS Pathogens 7.

Bressano M, Lorena Giachero M, Luna CM, Ducasse D a., 2010. An in vitro method for examining infection of soybean roots by Macrophomina phaseolina. Physiological and Molecular Plant Pathology 74, 201–204.

Brzostowski LF, Pruski TI, Hartman GL et al., 2018. Field evaluation of three sources of genetic resistance to sudden death syndrome of soybean. Theoretical and Applied Genetics, 1–12.

Cameron DD, Neal AL, van Wees SCMM, Ton J, 2013. Mycorrhiza-induced resistance: more than the sum of its parts? Trends in plant science 18, 539–45.

Chang H-X, Yendrek CR, Caetano-Anolles G, Hartman GL, 2016. Genomic characterization of plant cell wall degrading enzymes and in silico analysis of xylanses and polygalacturonases of Fusarium virguliforme. BMC Microbiology 16, 147.

Cheynier V, Comte G, Davies KM, Lattanzio V, Martens S, 2013. Plant phenolics: Recent advances on their biosynthesis, genetics, andecophysiology. Plant Physiology and Biochemistry 72, 1–20.

Conrath U, 2006. Systemic acquired resistance. Plant Signal Behav 1, 179–184.

Conrath U, Beckers GJM, Langenbach CJG, Jaskiewicz MR, 2015. Priming for enhanced defense. Annual review of phytopathology 53, 97–119.

Conrath U, Pieterse CMJ, Mauch-mani B, Mauch-mani B, 2002. Priming in plant – pathogen interactions. Trends in Plant Science 7, 210–216.

Cranenbrouck S, Voets L, Bivort C, Renard L, Strullu D-G, Declerck S, 2005. Methodologies for in vitro cultivation of arbuscular mycorrhizal fungi with root organs. In: In vitro culture of mycorrhizas. Springer, 341–375.

Crouzet J, Roland J, Peeters E et al., 2013. NtPDR1, a plasma membrane ABC transporter from Nicotiana tabacum, is involved in diterpene transport. Plant Molecular Biology 82, 181–192.

Dar MH, Reshi ZA, 2017. Vesicular Arbuscular Mycorrhizal (VAM) fungi-as a major biocontrol agent in modern sustainable agriculture system. Russian Agricultural Sciences 43, 138–143.

Declerck S, Strullu DG, Plenchette C, 1998. Monoxenic culture of the intraradical forms of Glomus sp. isolated from a tropical ecosystem: a proposed methodology for germplasm collection. Mycologia, 579–585.

Duhamel M, Vandenkoornhuyse P, 2013. Sustainable agriculture: possible trajectories from mutualistic symbiosis and plant neodomestication. Trends in plant science 18, 597–600.

Dunham WC, DunhamTrimmer LLC, 2015. Evolution and future of biocontrol. In: 10th Annual Biocontrol Industry Meeting (ABIM), Basel, Switzerland, October 20th.

Eke P, Chatue Chatue G, Wakam LN, Kouipou RMT, Fokou PVT, Boyom FF, 2016. Mycorrhiza consortia suppress the fusarium root rot (Fusarium solani f. sp. Phaseoli) in common bean (Phaseolus vulgaris L.). Biological Control 103, 240–250.

FAOSTAT, 2018. Food and Agriculture Organization of the United Nations.

Frenzel A, Tiller N, Hause B, Krajinski F, 2006. The conserved arbuscular mycorrhiza-specific transcription of the secretory lectin MtLec5 is mediated by a short upstream sequence containing specific protein binding sites. Planta 224, 792–800.

Gallou A, Declerck S, Cranenbrouck S, 2012. Transcriptional regulation of defence genes and involvement of the WRKY transcription factor in arbuscular mycorrhizal potato root colonization. Functional & integrative genomics 12, 183–98.

Gallou A, De Jaeger N, Cranenbrouck S, Declerck S, 2010. Fast track in vitro mycorrhization of potato plantlets allow studies on gene expression dynamics. Mycorrhiza 20, 201–207.

Gallou A, Lucero Mosquera HPHP, Cranenbrouck S et al., 2011. Mycorrhiza induced resistance in potato plantlets challenged by Phytophthora infestans. Physiological and Molecular Plant Pathology 76, 20–26.

Garcia-Garrido JM, Ocampo J a, García-Garrido JM, Ocampo J a, 2002. Regulation of the plant defence response in arbuscular mycorrhizal symbiosis. Journal of experimental botany 53, 1377–86.

Giachero ML, Marquez N, Gallou A, Luna CM, Declerck S, Ducasse D, 2017. An in vitro method for studying the three-way interaction between soybean, Rhizophagus irregularis and the soil-borne pathogen Fusarium virguliforme. 8, 1–9.

Hartman GL, Chang HX, Leandro LF, 2015. Research advances and management of soybean sudden death syndrome. Crop Protection 73, 60–66.

Hartman GL, Rupe JC, Sikora EF, Domier LL, Davis JA, Steffey KL, 2016. Compendium of Soybean Diseases and Pests, Fifth Edition (GL Hartman, JC Rupe, EJ Sikora, LL Domier, JA Davis, KL Steffey, Eds,). The American Phytopathological Society.

Ho VSM, Wong JH, Ng TB, 2007. A thaumatin-like antifungal protein from the emperor banana. Peptides 28, 760–766.

Iqbal MJ, Yaegashi S, Ahsan R, Shopinski KL, Lightfoot DA, 2005. Root response to Fusarium solani f. sp. glycines: temporal accumulation of transcripts in partially resistant and susceptible soybean. Theor Appl Genet 110, 1429–1438.

Jung SC, Martinez-Medina A, Lopez-Raez J a, Pozo MJ, 2012. Mycorrhiza-Induced Resistance and Priming of Plant Defenses. Journal of chemical ecology 38, 651–64.

Kandel YR, Wise KA, Bradley CA, Chilvers MI, Tenuta AU, Mueller DS, 2016. Fungicide and Cultivar Effects on Sudden Death Syndrome and Yield of Soybean. Plant Disease 100, 1339–1350.

Kumar A, Verma JP, 2018. Does plant—Microbe interaction confer stress tolerance in plants: A review? Microbiological Research 207, 41–52.

Lenoir I, Fontaine J, Lounès-Hadj Sahraoui A, 2016. Arbuscular mycorrhizal fungal responses to abiotic stresses: A review. Phytochemistry 123, 4–15.

Liu J, Maldonado-Mendoza I, Lopez-Meyer M, Cheung F, Town CD, Harrison MJ, 2007. Arbuscular mycorrhizal symbiosis is accompanied by local and systemic alterations in gene expression and an increase in disease resistance in the shoots. The Plant Journal 50, 529–544.

Liu J-J, Sturrock R, Ekramoddoullah AKM, 2010. The superfamily of thaumatin-like proteins: its origin, evolution, and expression towards biological function. Plant cell reports 29, 419–36.

López-Ráez J a, Verhage A, Fernández I et al., 2010. Hormonal and transcriptional profiles highlight common and differential host responses to arbuscular mycorrhizal fungi and the regulation of the oxylipin pathway. Journal of experimental botany 61, 2589–601.

Marquez N, Giachero ML, Gallou A et al., 2018. Transcriptional Changes in Mycorrhizal and Nonmycorrhizal Soybean Plants upon Infection with the Fungal Pathogen Macrophomina phaseolina. Molecular Plant-Microbe Interactions 31, 842–855.

McGonigle TP, Miller MH, Evans DG, Fairchild GL, Swan JA, 1990. A new method which gives an objective measure of colonization of roots by vesicular arbuscular mycorrhizal fungi. New phytologist 115, 495–501.

Migocka M, Papierniak A, Rajsz A, 2017. Cucumber PDR8/ABCG36 and PDR12/ABCG40 plasma membrane proteins and their up-regulation under abiotic stresses. Biologia Plantarum 61, 115–126.

Nadeem SM, Ahmad M, Zahir ZA, Javaid A, Ashraf M, 2014. The role of mycorrhizae and plant growth promoting rhizobacteria (PGPR) in improving crop productivity under stressful environments. Biotechnology Advances 32, 429–448.

Ngaki MN, Wang B, Sahu BB et al., 2016. Tanscriptomic study of the soybean-Fusarium virguliforme interaction revealed a novel ankyrin-repeat containing defense gene, expression of whose during infection led to enhanced resistance to the fungal pathogen in transgenic soybean plants. PLoS ONE 11.

Nivedita, Verma PK, Upadhyaya KC, 2017. Lectin Protein Kinase Is Induced in Plant Roots in Response to the Endophytic Fungus, Piriformospora indica. Plant Molecular Biology Reporter 35, 323–332.

Nuruzzaman M, Zhang R, Cao HZ, Luo ZY, 2014. Plant pleiotropic drug resistance transporters: Transport mechanism, gene expression, and function. Journal of Integrative Plant Biology 56, 729–740.

Olawuyi OJ, Odebode AC, Oyewole IO, Akanmu AO, Afolabi O, 2014. Effect of arbuscular mycorrhizal fungi on Pythium aphanidermatum causing foot rot disease on pawpaw (Carica papaya L.) seedlings. Archives of Phytopathology and Plant Protection 47, 185–193.

Oyewole BO, Olawuyi OJ, Odebode AC, Abiala MA, 2017. Influence of Arbuscular mycorrhiza fungi (AMF) on drought tolerance and charcoal rot disease of cowpea. Biotechnology Reports 14, 8–15.

Pandin C, Le Coq D, Canette A, Aymerich S, Briandet R, 2017. Should the biofilm mode of life be taken into consideration for microbial biocontrol agents? Microbial Biotechnology 10, 719–734.

Phillips JM, Hayman DS, 1970. Improved procedures for clearing roots and staining parasitic and VA mycorrhizal fungi for rapid assessment of infection. Transactions of the British Mycological Society 55, 158–161.

Pieterse CMJ, Zamioudis C, Berendsen RL, Weller DM, Van Wees SCM, Bakker PAHM, 2014. Induced Systemic Resistance by Beneficial Microbes. Annual Review of Phytopathology 52, 347–375.

Pozo MJ, Azcón-Aguilar C, 2007. Unraveling mycorrhiza-induced resistance. Current opinion in plant biology 10, 393–8.

Pozo MJ, Verhage A, García-andrade J, García JM, Azcón-aguilar C, 2009. Priming Plant Defence Against Pathogens by Arbuscular Mycorrhizal Fungi., 1–13.

Radwan O, Liu Y, Clough SJ, 2011. Transcriptional analysis of soybean root response to Fusarium virguliforme, the causal agent of sudden death syndrome. Molecular plant-microbe interactions 24, 958–972.

Rather IA, Awasthi P, Mahajan V, Bedi YS, Vishwakarma RA, Gandhi SG, 2015. Molecular cloning and functional characterization of an antifungal PR-5 protein from Ocimum basilicum. Gene 558, 143–151.

Di Rienzo JA, Casanoves F, Balzarini MG, Gonzalez L, Tablada M, Robledo CW, 2013. INFOSTAT.

Di Rienzo JA, Guzmán AW, Casanoves F, 2002. A multiple-comparisons method based on the distribution of the root node distance of a binary tree. Journal of agricultural, biological, and environmental statistics 7, 129–142.

Roy KW, State M, Rupe JC et al., 1997. Sudden Death Syndrome of Soybean. Plant Disease 81, 1100–1111.

S Navi S, Rajasab A, 2016. In Vitro Evaluation of Commercial Fungicides against some of the Major Soil Borne Pathogens of Soybean. Journal of Plant Pathology & Microbiology 07.

Sahu BB, Baumbach JL, Singh P, Srivastava SK, Yi X, Bhattacharyya MK, 2017. Investigation of the fusarium virguliforme transcriptomes induced during infection of soybean roots suggests that enzymes with hydrolytic activities could play a major role in root necrosis. PLoS ONE 12, 1–22.

Schouteden N, Waele D De, Panis B, Vos CM, 2015. Arbuscular mycorrhizal fungi for the biocontrol of plant-parasitic nematodes: A review of the mechanisms involved. Frontiers in Microbiology 6, 1–12.

Schubler A, 2010. The Glomeromycota: a species list with new families and new genera. http://www.amf-phylogeny.com.

Shigeto J, Tsutsumi Y, 2016. Diverse functions and reactions of class III peroxidases. New Phytologist 209, 1395–1402.

Siah A, Magnin-robert M, Randoux B, Choma C, Halama P, Reignault P, 2018. Natural Agents Inducing Plant Resistance Against Pests and Diseases.

Singh NK, Kumar KRR, Kumar D, Shukla P, Kirti PB, 2013. Characterization of a Pathogen Induced Thaumatin-Like Protein Gene AdTLP from Arachis diogoi, a Wild Peanut (J-H Liu, Ed,). PLoS ONE 8.

Singh P, Kuo Y-C, Mishra S et al., 2012. The Lectin Receptor Kinase-VI.2 Is Required for Priming and Positively Regulates Arabidopsis Pattern-Triggered Immunity. The Plant Cell 24, 1256–1270.

Smith SE, Read DJ, 2008. Mycorrhizal symbiosis. 3rd. Academic Press New York, ISBN 440026354, 605.

Srivastava SK, Huang X, Brar HK, Fakhoury AM, Bluhm BH, Bhattacharyya MK, 2014. The genome sequence of the fungal pathogen Fusarium virguliforme that causes sudden death syndrome in soybean. PLoS ONE 9.

Srour AY, Gibson DJ, Leandro LFS, Malvick DK, Bond JP, Fakhoury AM, 2017. Unraveling Microbial and Edaphic Factors Affecting the Development of Sudden Death Syndrome in Soybean. Phytobiomes 1, 91–101.

Vigers AJ, Wiedemann S, Roberts WK, Legrand M, Selitrennikoff CP, Fritig B, 1992. Thaumatin-like pathogenesis-related proteins are antifungal. Plant Science 83, 155–161.

Voets L, de la Providencia IE, Fernandez K, Ijdo M, Cranenbrouck S, Declerck S, 2009. Extraradical mycelium network of arbuscular mycorrhizal fungi allows fast colonization of seedlings under in-vitro conditions. Mycorrhiza 19, 347–356.

Wang Y, Bouwmeester K, 2017. L-type lectin receptor kinases: New forces in plant immunity. PLoS Pathogens 13, 1–7.

Wang B, Qiu YL, 2006. Phylogenetic distribution and evolution of mycorrhizas in land plants. MYCORRHIZA 16, 299–363.

Wen Z, Tan R, Yuan J et al., 2014. Genome-wide association mapping of quantitative resistance to sudden death syndrome in soybean. BMC Genomics 15, 1–11.

Woo JY, Jeong KJ, Kim YJ, Paek K-H, 2016. *CaLecRK-S.5*, a pepper L-type lectin receptor kinase gene, confers broad-spectrum resistance by activating priming. Journal of Experimental Botany, erw336.

Wrather A, Shannon G, Balardin R et al., 2010. Effect of diseases on soybean yield in the top eight producing countries in 2006. Plant Health Progress doi 10, 2008–2013.

Xu Z, Song N, Ma L, Fang D, Wu J, 2018. Plant Diversity NaPDR1 and NaPDR1-like are essential for the resistance of Nicotiana attenuata against fungal pathogen Alternaria alternata. Plant Diversity 40, 68–73.

Yuan J, Zhu M, Lightfoot DA, Javed MJ, Yang JY, Meksem K, 2008. In silico comparison of transcript abundances during Arabidopsis thaliana and Glycine max resistance to Fusarium virguliforme. BMC Genomics 9.

Zhang Q, Gao X, Ren Y et al., 2018. Improvement of Verticillium wilt resistance by applying arbuscular mycorrhizal fungi to a cotton variety with high symbiotic efficiency under field conditions. International Journal of Molecular Sciences 19.

